# Development and Mobile Deployment of a Stair Recognition System for Human-Robot Locomotion

**DOI:** 10.1101/2023.04.25.538248

**Authors:** Andrew Garrett Kurbis, Alex Mihailidis, Brokoslaw Laschowski

**Affiliations:** Department of Systems Design Engineering, University of Waterloo, Waterloo, ON Canada; Temerty Faculty of Medicine, University of Toronto, Toronto, ON Canada; Toronto Rehabilitation Institute, University Health Network, Toronto, ON Canada; Department of Mechanical and Industrial Engineering, University of Toronto, Toronto, ON Canada

**Keywords:** computer vision, deep learning, exoskeletons, prosthetics, wearable robotics

## Abstract

Environment sensing and recognition can improve the safety and autonomy of human-robot locomotion, especially during transitions between environmental states such as walking to and from stairs. However, accurate and real-time perception on edge devices with limited computational resources is an open problem. Here we present the development and mobile deployment of *StairNet* - a vision-based automated stair recognition system powered by deep learning. Building on *ExoNet* - the largest open-source dataset of egocentric images of real-world walking environments - we designed a new dataset specifically for stair recognition with over 515,000 images. We then developed a lightweight and efficient convolutional neural network for image classification, which accurately predicted complex stair environments with 98.4% accuracy. We also studied different model compression and optimization methods and deployed our system on several mobile devices running a custom-designed iOS application with onboard accelerators using CPU, GPU, and/or NPU backend computing. Of the designs that we tested, our highest performing system showed negligible reductions in classification accuracy due to the model conversion for mobile deployment and achieved an inference time of 2.75 ms on an iPhone 11. The high speed and accuracy of the *StairNet* system on edge devices opens new opportunities for autonomous control and planning of robotic prosthetic legs, exoskeletons, and other assistive technologies for human locomotion.

## I. INTRODUCTION

HUMANS rely on a biological system feedback loop for legged locomotion [1], [2]. This feedback loop includes: 1) recognition of the walking environment using the human visual system, 2) cognitive processing of the environment state and locomotor intent through neural control, 3) translation of the locomotor intent into movement through the musculoskeletal system, and 4) the physical environment response (i.e., the new walking environment that the human interacts with). Disruptions to this control loop can occur from limitations to the musculoskeletal system due to aging and/or physical disabilities such as amputation, or through communication limitations to the central nervous system due to stroke or spinal cord injury, all of which can affect one’s ability to perform locomotor activities and navigate new environments safely and effectively [3].

Similar to the biological vision-locomotor feedback loop, autonomous control of robotic leg prostheses and exoskeletons requires continuous assessment of the environmental states for seamless transitions between different locomotion modes, each of which have individually tuned control parameters [4]. Accurate classification of stairs can be particularly important due to safety implications and the greater risk of severe injuries if the environment is misclassified during stair locomotion. However, robotic prostheses and exoskeletons typically rely on mechanical, inertial, and/or electromyographic (EMG) sensors for state estimation, which are generally limited to the current state of the human-robot-environment system, which is analogous to walking blindfolded. Computer vision can allow for environment-adaptive control via forward prediction of the environmental states in order to assist transitions to and from stairs (Fig. 1).

**Fig. 1.**
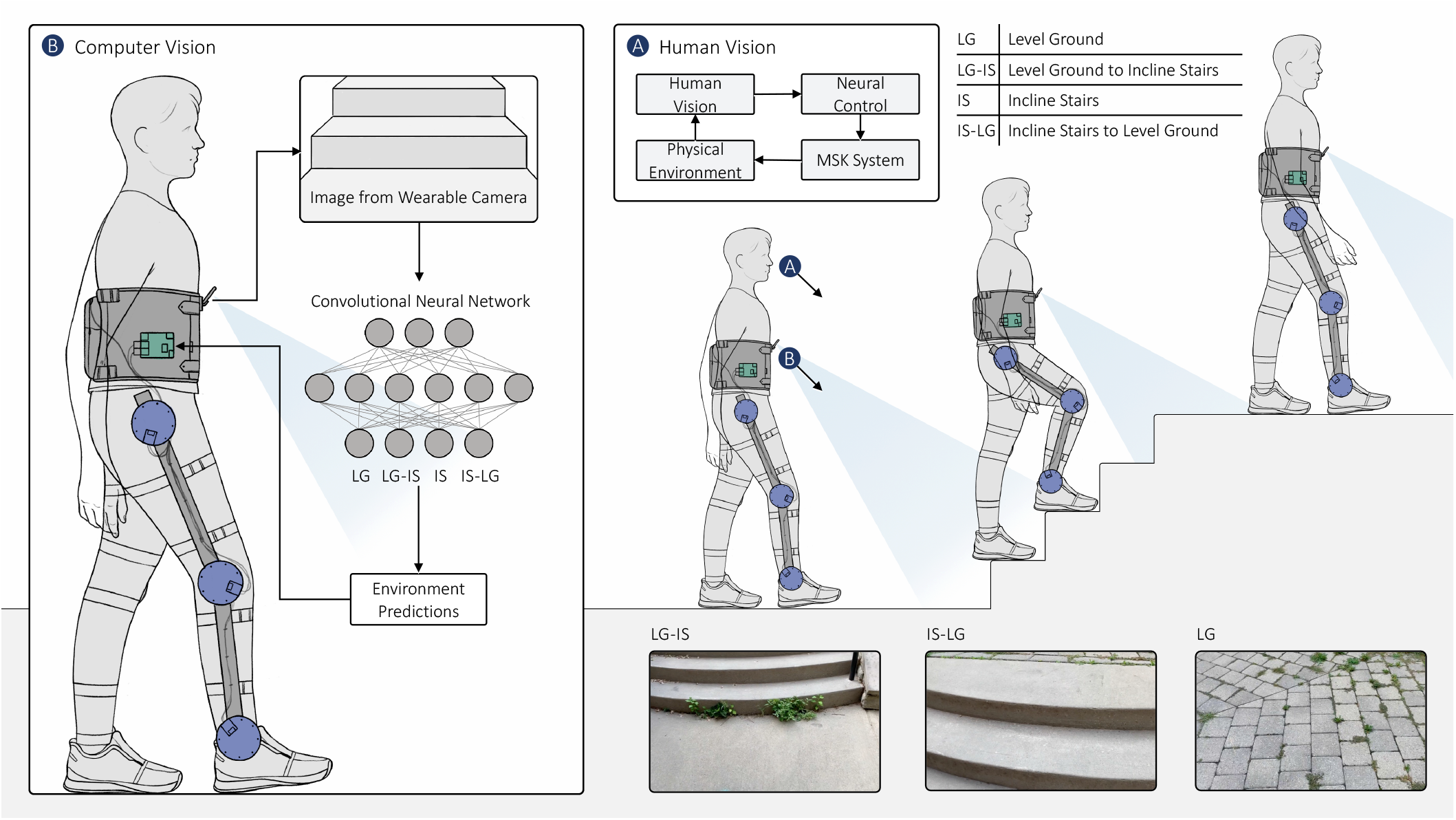
Schematic showing the application of computer vision and deep learning for sensing and classification of real-world stair environments during human-robot locomotion.

Although vision-based stair recognition systems have been developed for wearable robotics [5]–[13], these systems have used statistical pattern recognition and machine learning algorithms that require manual feature engineering and/or have involved relatively small datasets, which can limit their generalizability to diverse walking environments [14]. While some researchers have explored using large-scale deep learning for robust perception, these systems have mainly been evaluated from a theoretical performance standpoint on testing sets. For example, we recently presented the theoretical framework for a vision-based stair recognition system powered by deep learning [15] and used cloud computing for the development and testing.

It remains an open question how best to develop perception systems that are accurate and fast on edge devices with limited computational and memory resources. Convolutional neural networks (CNNs) with deeper layers tend to be more accurate for image classification, with the tradeoff of more parameters and operations, which can affect prediction speed [16]. This research question is especially pertinent to human-robot locomotion, where accuracy is important for user safety, while inference speed is important for real-time control and adapting online to different walking environments. Recent advances in lightweight and efficient deep learning models such as MobileNets [17], [18], and developments in AI accelerators like neural processing units (NPUs) [19], are allowing perception systems to be deployed on mobile and embedded computing devices.

Building on our previous theoretical research [15], here we studied the effects of different parameters on prediction speed and accuracy when deployed on mobile devices in order to optimize the system for real-world applications. We studied different deep learning model and training parameters such as learning rate, batch size, dropout rate, and number of frozen layers in transfer learning, as well as different mobile devices (i.e., iPhone 8+, iPhone X, iPhone 11, and iPhone 13), model compression and optimization methods for deployment, and backend processing options such as single-threaded CPU, multi-threaded CPU, GPU, and a combination of CPU, GPU, and NPU in parallel.

This study resulted in the development of *StairNet* - the largest and most robust AI-powered stair recognition system developed to date with over 98% accuracy and the first to be deployed on mobile devices and show real-time inference predictions, with speeds up to 2.8 ms. In light of these results, the *StairNet* system has the ability to support the development of next-generation autonomous controllers for robotic prosthetic legs, exoskeletons, and other assistive technologies for human locomotion.

## II. Methods

### A. Computer Vision Dataset

In order to study and optimize our perception system for real-world applications, we developed a new image dataset us-ing manually annotated images from *ExoNet* [20] – the largest and most diverse open-source dataset of wearable camera images of walking environments. Images from six of the twelve original *ExoNet* classes were used, which included environments that a user would encounter during stair ascent. The initial dataset included ∼543,000 RGB images. We then reduced the number of classes in the dataset from 6 to 4 by grouping two classes initially separated in *ExoNet* based on the presence of doors/walls in the “level ground” and “incline stairs” classes. The final four environment classes included: 1) level ground (LG), which included both *ExoNet* classes “level ground steady state” and “level ground transition to door/wall,” 2) level ground – incline stairs (LG-IS), which consisted of the *ExoNet* class “level ground transition to incline stairs,” 3) incline stairs (IS), which included both *ExoNet* classes “incline stairs steady state” and “incline stairs transition to door/wall,” and 4) incline stairs – level ground (IS-LG), which consisted of the *ExoNet* class “incline stairs transition to level ground.” The dataset was randomly split into training (89.5%), validation (3.5%), and test (7%) sets, matching the subset distribution values from the original *ExoNet* studies [21]-[23]. Each subset maintained the overall class distributions (i.e., 85.8% for LG, 9.3% for IS, 3.1% for LG-IS, and 1.8% for IS-LG).

### B. Convolutional Neural Network

We then developed a convolutional neural network using the base model of MobileNetV2 for image classification [17], [18] (Fig. 2), which was designed by Google for mobile and embedded vision applications with an architecture based on depthwise separable convolutions. The model uses width and resolution multipliers to create a lightweight framework with the trade-off of slightly lower accuracy for significantly reduced computational requirements. Its efficient and lightweight design can allow for onboard real-time computations for robot control. We developed the deep learning model in TensorFlow 2.7 [24]. We studied the effects of different model parameters using a Google Cloud Tensor Processing Unit (TPU), a hardware accelerator optimized for efficient machine learning via large matrix operations. An initial model was developed using the default parameter values from Laschowski et al. [21]-[23], who trained MobileNetV2 and a dozen other state-of-the-art deep learning models on the whole *ExoNet* database. Our initial model included the base model of MobileNetV2, randomly initialized weights, ∼2.3 million parameters, a batch size of 128, and a base learning rate of 0.001. The model was trained using the Adam optimizer [25] and a cosine weight decay learning rate policy.

**Fig. 2.**
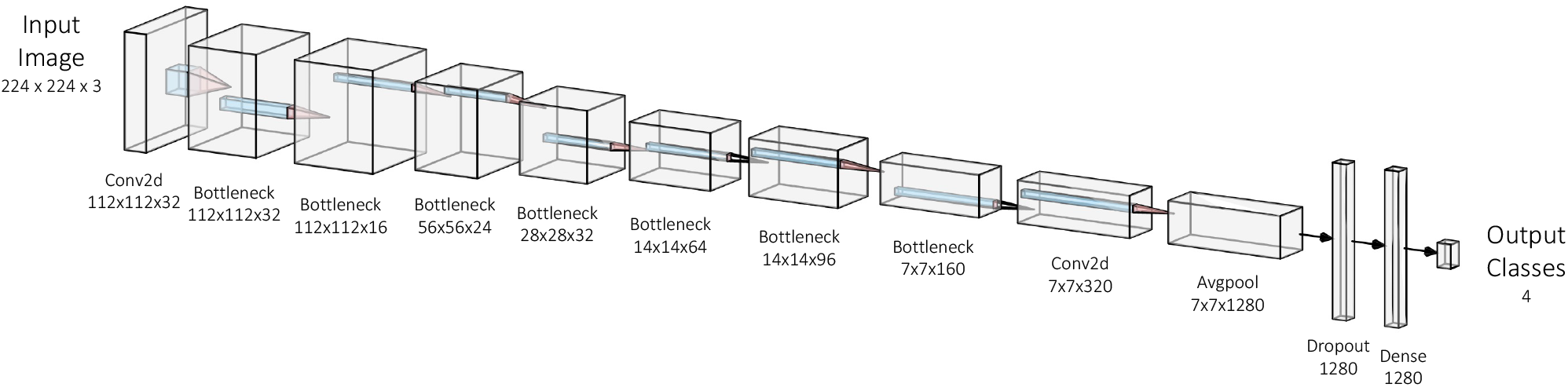
The MobileNetV2 neural network architecture [17] used for image classification of stair environments. The network has ∼2.3 million parameters and 0.6 GFLOPs.

### C. StairNet Dataset

Our initial study on the training and validation sets using the preliminary neural network revealed low categorical accuracies of 53.6% and 66.4% in the transition classes IS-LG and LG-IS, respectively. We investigated the cause of this low accuracy as the performance was significantly lower than other stair recognition systems trained on smaller datasets, such as Laschowski and colleagues [6] who achieved 94.9% prediction accuracy on ∼34,000 images. Our analysis revealed that several images in the *ExoNet* database were misclassified or had significant camera obstructions (e.g., the lens covered by a hand) where the assessment of the walking environments was not possible. These images were found throughout the training, validation, and testing sets, thereby impacting both the development and evaluation of our preliminary deep learning model.

To fix these errors, we manually re-labelled the images. We proposed and developed new class definitions to increase the precision of the cut-off points between different environment classes (Table 1). After three manual annotation passes of the dataset, images that were considered out of scope (e.g., an image of a wall without level ground visible) and images that had significant camera obstructions were discarded, therein reducing the total number of images in our new dataset to ∼515,000. We then randomly split the dataset into training (89.5%), validation (3.5%), and test (7%) sets while maintaining the class distributions (Table 2). After completing the dataset revision, our preliminary deep learning model was retested on the new validation set with the same parameters as previously used. The categorical accuracies improved from 53.6% to 84.4% for the IS-LG class and from 66.4% to 87.5% for the LG-IS class. These results showed that our new class definitions allow for more precise cut-off points between the different environment classes. Our new image dataset was uploaded to IEEE DataPort and is publicly available to the research community for download https://ieeedataport.org/documents/stairnet-computer-vision-dataset-stairrecognition.

**Table 1.** The new class definitions that we developed and used to manually label our computer vision dataset called *StairNet*.

| Class | Class description |
| --- | --- |
| LG | An image that contains a level ground environment where incline stairs are not clearly visible. |
| LG-IS | An image with incline stairs where the horizontal surface area of the bottom step or landing is clearly greater than the surface area of other steps visible in the image (i.e., the surface area or depth is approximately 1.5x the size of subsequent steps). |
| IS | An image with multiple incline stairs where the horizontal surface area of the top and bottom step or landing is not clearly greater than one another. |
| IS-LG | An image with incline stairs where the horizontal surface area of the top step or landing is clearly greater than that of other steps or landings visible in the image (i.e., the surface area or depth is approximately 1.5x the size of subsequent steps). For an incline stair to be included in the IS-LG class, the horizontal face of the last step prior to level ground must be visible. |

**Table 2.** Normalized confusion matrix of the environment predictions on the *StairNet* validation set using the oversampled model with a minimum value of 25,000 images per class. The horizontal and vertical axes are the predicted and labelled classes, respectively.

|  |  |  |  |  |
| --- | --- | --- | --- | --- |
| IS | 96.3 | 1.8 | 0.4 | 1.5 |
| IS-LG | 6.4 | 86.0 | 6.7 | 1.0 |
| LG | 0.1 | 0.1 | 99.6 | 0.3 |
| LG-IS | 3.2 | 0.0 | 3.8 | 92.3 |
|  | IS | IS-LG | LG | LG-IS |

The images that were discarded from the training set were considered out of scope such that they did not contain either level-ground terrain or incline stairs. As the *StairNet* system is designed specifically for stair recognition, there is no loss of characteristics related to the intended application as any classifications made outside of the *StairNet* classes are considered out of scope and thus require additional models for image classification.

### D. Model Optimization

We then studied the effects of different model parameters on performance using our new *StairNet* dataset to optimize the validation accuracy and loss. We first studied the use of transfer learning with pre-trained weights from ImageNet [26]. Transfer learning and the addition of a global average pooling 2D layer and a softmax dense prediction layer decreased the number of trainable parameters, thus reducing the time and computational requirements to optimize the model. We studied five freeze layer parameter values to evaluate the use of transfer learning at 141, 100, 50, 25, and 5 frozen layers. Each model variation was run for 60 epochs. The optimal freeze layer value was 5 frozen layers with ∼2.2 million parameters, which resulted in the highest validation accuracy and lowest validation loss.

We also studied the effects of different batch sizes (64, 128, 256) and base learning rates (0.0001, 0.00001, 0.000001) on the validation accuracy and loss. Each parameter combination was run for 60 epochs. The optimal combination was a batch size of 256 and a base learning rate of 0.0001. We then studied and compared the optimized pre-trained model to an equivalent non-pretrained model with randomly initialized weights. After 60 epochs, the accuracy of both models plateaued. The pre-trained model had a higher validation accuracy (∼98%) compared to the non-pretrained model (∼97%) while having similar final validation loss values. After 60 epochs, the pretrained model exhibited characteristics of overfitting such that an increase in the validation loss was observed. We studied the effects of an additional dropout layer, with dropout rates of 0.01, 0.02, and 0.05. A dropout rate of 0.02 resulted in the highest validation accuracy and lowest validation loss. We also studied the effects of L2 weight regularization. The models were run for 60 epochs, and the validation loss and accuracy were compared, which showed no additional performance benefit from the weight regularization.

To reduce overfitting, we studied the effects of oversampling of the underrepresented classes, specifically IS-LG and LG-IS. We randomly resampled and augmented images that were previously presented to the model during training. Five model designs were tested, which included different values for the minimum number of images per class required for model training (i.e., 25,000, 40,000, 60,000, 200,000, and 400,000).

The overall validation accuracy decreased as the minimum value increased. However, the categorical accuracy for the underrepresented classes increased as the minimum value increased, creating a more even categorical accuracy distribution across the different walking environments (Tables 3 and 4). Given that more significant consequences could result from a false negative than a false positive in the detection of stairs for robot control, we decided to select the deep learning model that oversampled with a minimum value of 400,000 images per class as this model had a more even categorical accuracy distribution and the lowest probability of false negatives as seen in the reduced probability of misclassification as LG in class IS (0.3%) and IS-LG (2.2%).

**Table 3.** Normalized confusion matrix of the environment predictions on the *StairNet* validation set using the oversampled model with a minimum value of 400,000 images per class. The horizontal and vertical axes are the predicted and labelled classes, respectively.

|  |  |  |  |  |
| --- | --- | --- | --- | --- |
| IS | 96.6 | 1.3 | 0.3 | 1.8 |
| IS-LG | 5.7 | 90.8 | 1.9 | 1.6 |
| LG | 0.1 | 0.3 | 99.0 | 0.6 |
| LG-IS | 3.2 | 0.4 | 2.2 | 94.2 |
|  | IS | IS-LG | LG | LG-IS |

**Table 4.** Distribution of the environment classes in the *ExoNet* database [20] and our new dataset called *StairNet*. The number of images (#) and percent of the dataset (%) are reported for each class.

| Class | ExoNet (#) | ExoNet (%) | StairNet (#) | StairNet (%) |
| --- | --- | --- | --- | --- |
| LG | 456,775 | 84.14 | 442,360 | 85.82 |
| LG-IS | 26,067 | 4.80 | 15,888 | 3.08 |
| IS | 48,986 | 9.02 | 48,179 | 9.35 |
| IS-LG | 11,040 | 2.03 | 9,025 | 1.75 |

Finally, we fine-tuned the oversampled model with different batch sizes and base learning rates on a high epoch run to further increase the categorical and overall validation accuracies. By comparing the validation accuracy, loss, and confusion matrices, the final model was selected with a reduced base learning rate of 0.00001, a batch size of 128, and a cosine weight decay learning policy. The model also included pretrained weights, 5 frozen layers, ∼2.2 million parameters, a minimum categorical image count of 400,000 images, and an additional dropout layer with a dropout rate of 0.02. The final model was trained for 60 epochs (Fig. 3).

**Fig. 3.**
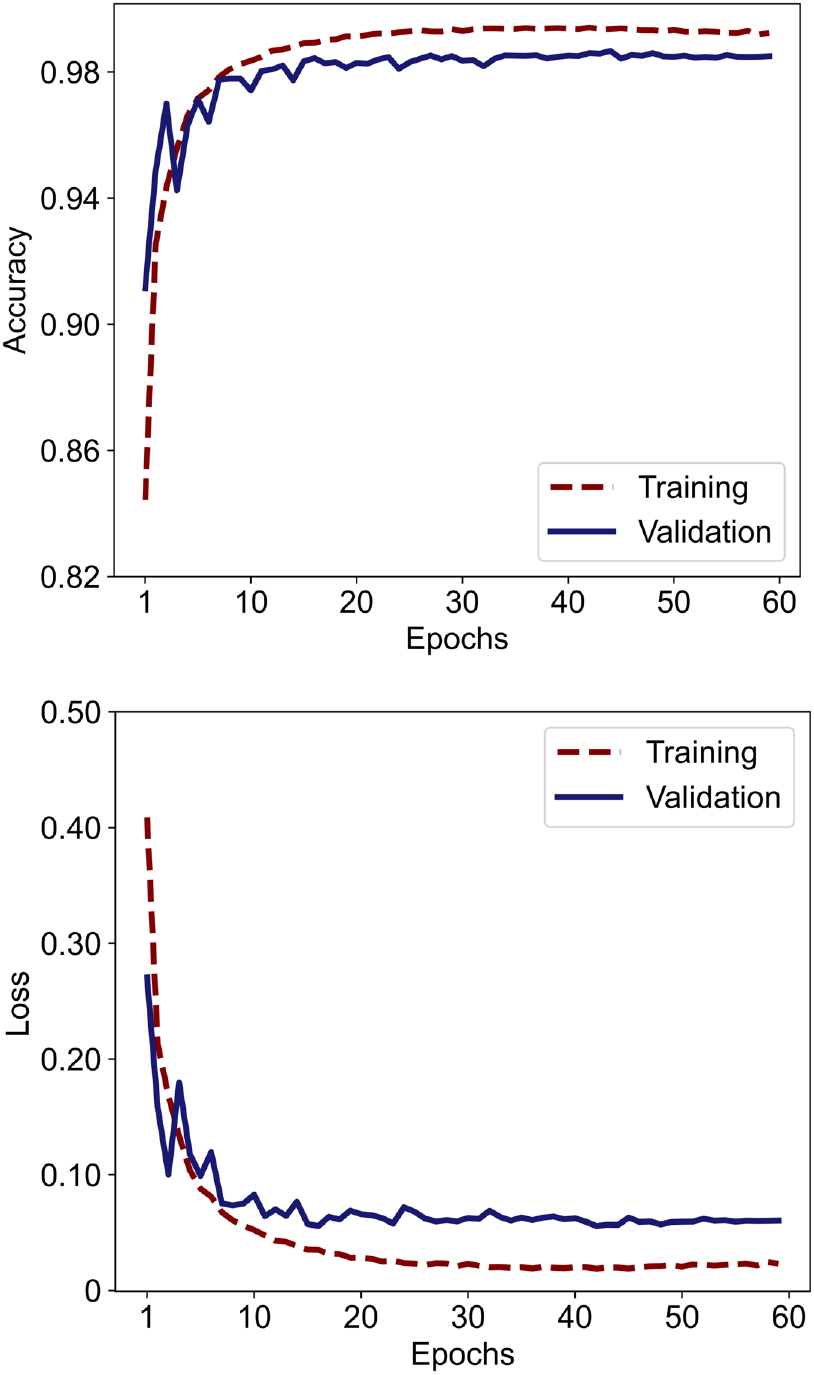
The image classification accuracy and loss on the *StairNet* training and validation sets using our optimized deep learning model.

### D. Mobile Deployment

To study and optimize the perception system for real-world applications, we used TensorFlow Lite (TFLite) for mobile deployment and inference (Fig. 4) [27]. TFLite has advantages over other forms of deployment like cloud computing due to its ability to run offline and perform inference on edge devices without internet connection requirements or the need to perform round trips to a machine learning server. Performing offline inference can reduce power requirements and privacy concerns as no data is required to leave the device, which can be especially important in clinical applications. TFLite also has a small binary size and supports highly efficient models, allowing for low inference times with minimal impact on accuracy during compression. We converted our deep learning model from its original h5 format into a TFLite flat buffer format compatible with the TFLite interpreter. The interpreter is the on-device infrastructure that allows for mobile inference and supports multiple backend processing options like central processing units (CPUs), graphics processing units (GPUs), and neural processing units (NPUs).

**Fig. 4.**
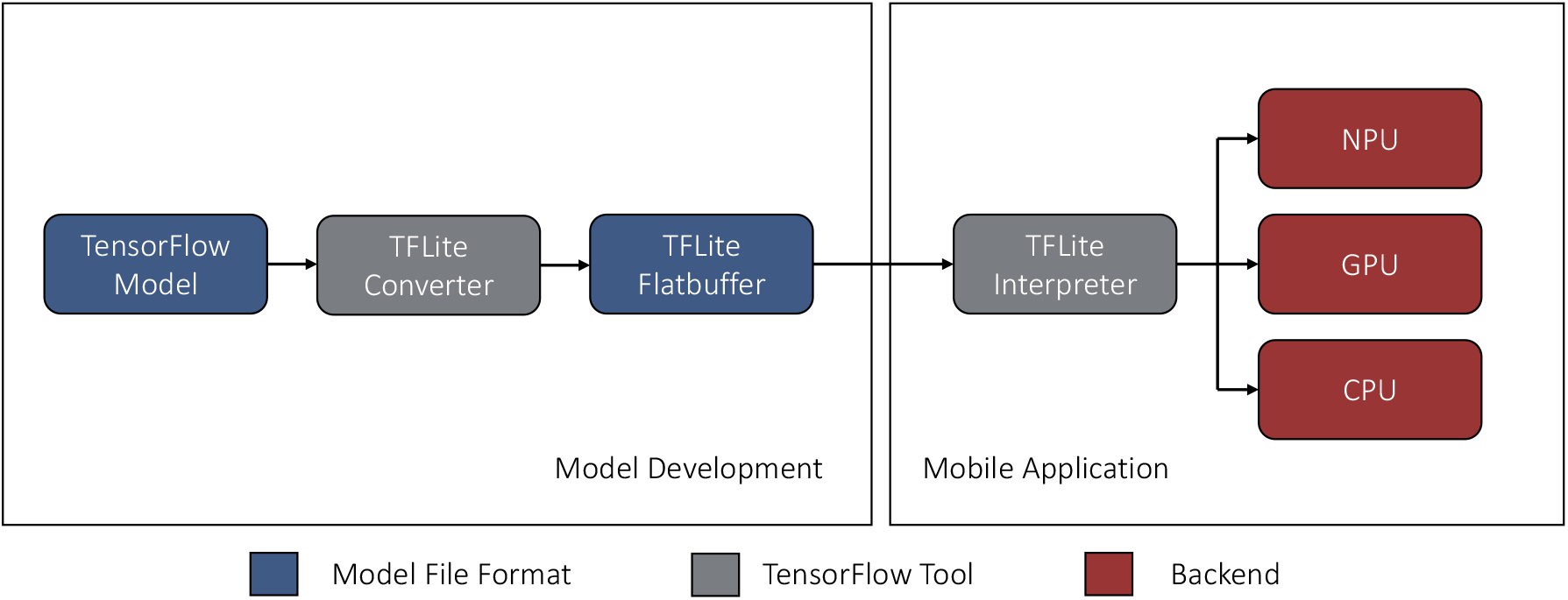
Conversion and mobile deployment of our deep learning model with different onboard computational delegates, including a central processing unit (CPU), graphics processing unit (GPU), and neural processing unit (NPU).

Model optimization can benefit mobile deployment in terms of model size reduction, latency reduction, and accelerator compatibility. However, conversion and optimization methods tend to have a slight trade-off in accuracy. We studied five different model compression and optimization methods with increasing levels of precision reduction to determine the optimal method for our computer vision model and application. These included 1) float32 compression for general TFLite deployment, 2) post-training float16 quantization to reduce the model size and increase performance on CPU and GPU hardware, 3) post-training int8 weight quantization to reduce the model size and increase performance on CPU hardware, 4) post-training quantization with int16 activations to reduce the model size and provide model compatibility with integer-only hardware accelerators, and 5) post-training int8 full model quantization (i.e., model weights, biases, and activations) to reduce the model size and increase processor compatibility (Table 5).

**Table 5.** The model conversion and optimization methods that we studied and their resulting image classification accuracies on the *StairNet* validation set.

| Method | Conversion method | Size (MB) | Accuracy (%) | Accuracy reduction (percentage points) |
| --- | --- | --- | --- | --- |
|  | **Original model** | **27.6 MB** | **98.41%** | **0.000** |
| 1 | TFLite float32 compression | 8.9 MB | 98.41% | 0.001 |
| 2 | Post-training float16 quantization | 4.5 MB | 98.41% | 0.003 |
| 3 | Post-training int8 weight quantization | 2.7 MB | 98.41% | 0.001 |
| 4 | Post-training int8 quantization with int16 activations | 2.5 MB | 98.31% | 0.100 |
| 5 | Post-training int8 full model quantization | 2.5 MB | 98.30% | 0.111 |

We developed an iOS application to deploy our model using Swift 5 and Xcode 13.4.1 [28] (Fig. 5). The input image is prepared by taking a square crop from the camera feed and scaled to match the input resolution of our model (224×224). The output tensor is converted into a float array to obtain the confidence values for each class, which are passed to the mobile user interface with the top three classes displayed based on the highest confidence values. The application also displays the onboard inference time (ms). The class with the highest confidence for each image is compared to the image labels from the test set to estimate the overall prediction accuracy. This accuracy is compared to the original value from the *StairNet* model to evaluate any changes in accuracy due to the model conversion for mobile deployment.

**Fig. 5.**
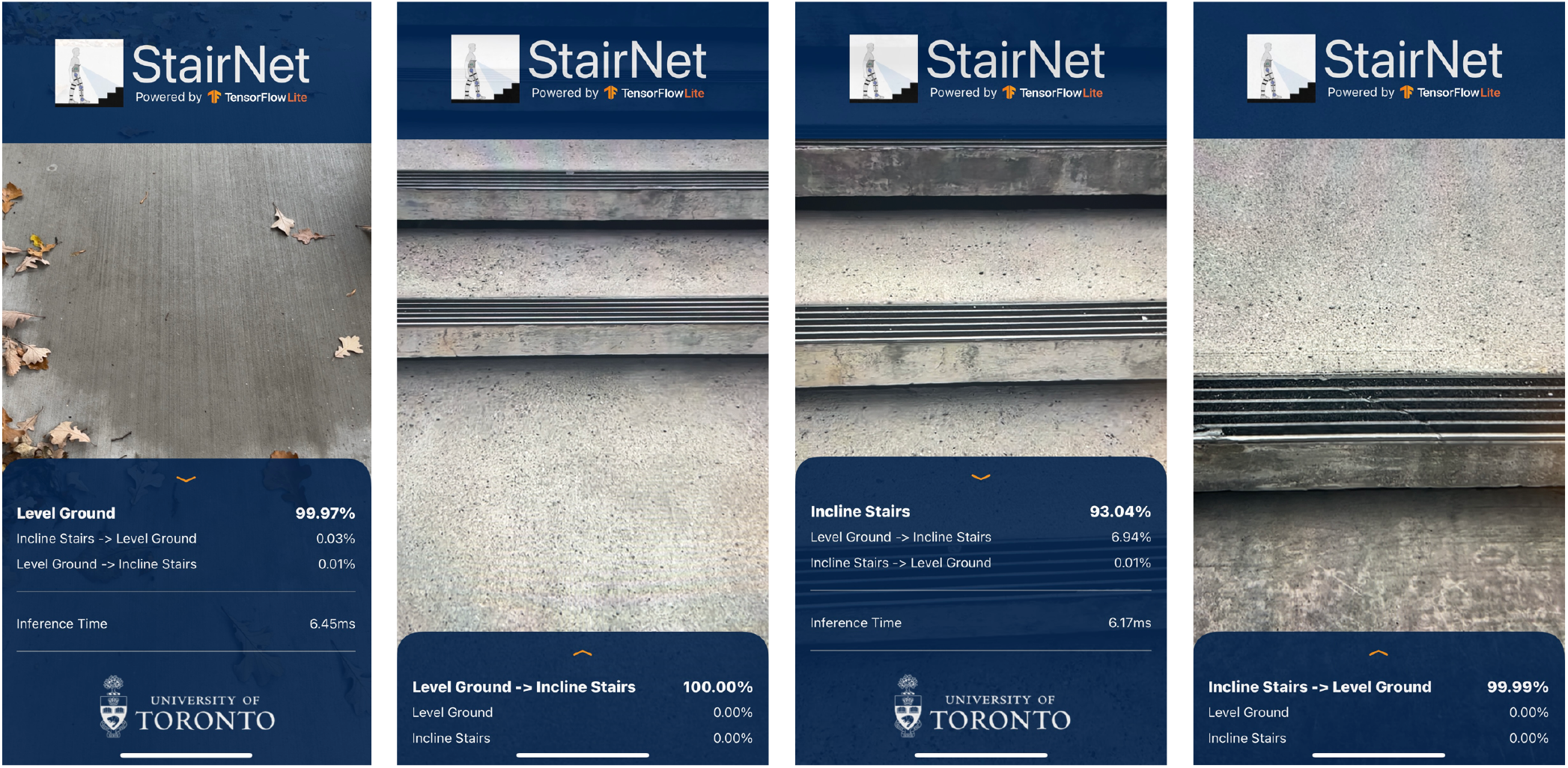
Our custom-designed iOS application for mobile deployment and inference of the *StairNet* automated stair recognition system.

To further study model compression and optimization methods, we tested each method on four different mobile devices (i.e., iPhone 8+, iPhone X, iPhone 11, and iPhone 13) with four different backend processing options, including a single-threaded CPU, a multi-threaded CPU, GPU, and a combination of CPU, GPU, and NPU in parallel. The backend processing options were created using previously developed APIs with access to hardware accelerators. The APIs included the Apple Metal delegate for direct GPU computing and the Apple CoreML delegate, which uses the three iOS processing options to maximize performance while minimizing memory usage and power consumption. Each device and backend processing option were tested and compared using a 90-second video consisting of stair ascent in an indoor and outdoor environment to best represent the performance of our system during real-world deployment. The average inference time was calculated by sampling at five-second intervals.

## III. Results

The stair recognition accuracies on the training and validation sets were 99.3% and 98.5%, respectively. When evaluated on the test set, our deep learning model achieved an overall image classification accuracy of 98.4%, a weighted f1 score of 98.4%, a weighted precision value of 98.5%, and a weighted recall value of 98.4%. Here, classification accuracy is defined as the number of images correctly identified by the neural network (35,507 images) out of the total number of images in the test set (36,085).

The classification accuracy on the test set varied between different environments, with a categorical accuracy of 99.0% for the LG class, 91.7% for the LG-IS class, 96.9% for the IS class, and 90.5% for the IS-LG class. Table 6 shows the normalized confusion matrix, which illustrates the classification performance distribution during inference. The two transition classes (i.e., LG-IS and IS-LG) achieved the lowest categorical accuracies, likely due to having the smallest class distributions, comprising only 3.1% and 1.8% of the total number of images in the dataset, respectively.

**Table 6.** Normalized confusion matrix of the environment predictions on the *StairNet* test set. The horizontal and vertical axes are the predicted and labelled classes, respectively.

|  |  |  |  |  |
| --- | --- | --- | --- | --- |
| IS | 96.9 | 1.5 | 0.3 | 1.3 |
| IS-LG | 6.5 | 90.5 | 2.4 | 0.6 |
| LG | 0.1 | 0.3 | 99.0 | 0.7 |
| LG-IS | 4.3 | 0.5 | 3.4 | 91.7 |
|  | IS | IS-LG | LG | LG-IS |

Fig. 6 shows examples of failure cases. The first row contains images from the LG class that were incorrectly classified as LGIS. The images contain features common within the LG-IS class, such as horizontal lines in the top section of the images, which were likely confused as transitions to incline stairs. The second row contains LG images that were incorrectly classified as IS. The images contain horizontal lines resulting from surface textures (e.g., brick flooring in the second column) that are present throughout the images and were likely confused for steady-state incline stairs. The bottom row contains images from the stair classes that were misclassified as LG. These false negatives pose a significant safety risk to users and contain unique stair characteristics, such as unusual materials (e.g., wood) or angles, which rotate the horizontal features to a diagonal or vertical axis.

**Fig. 6.**
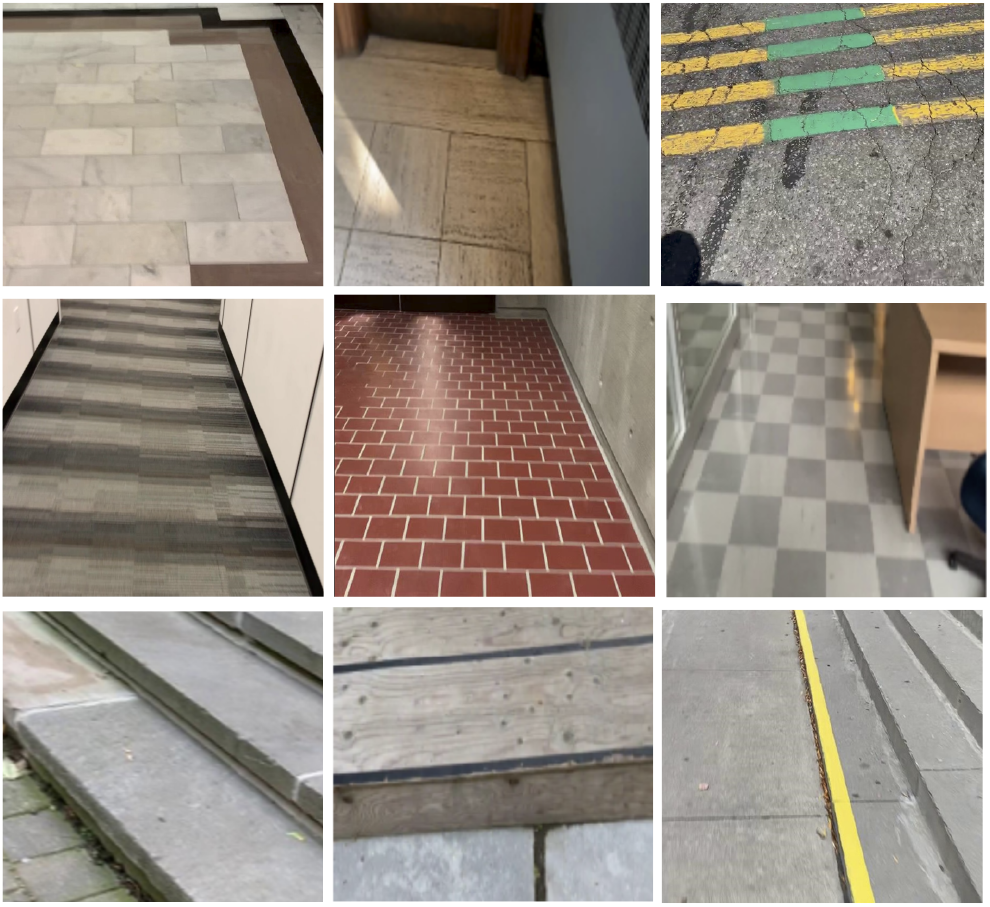
Examples of incorrect environment predictions by our deep learning model. The first row contains level ground images that were misclassified as level ground – incline stairs (LG-IS). The second row contains level ground images that were misclassified as incline stairs (IS). The third row contains images from the stair classes that were misclassified as level ground (LG).

The model compression and optimization methods that we studied only decreased the prediction accuracies between 0.001-0.111 percentage points compared to the original model. Float32 compression and float16 quantization maintained the greatest model precision and resulted in the highest post-conversion accuracy (98.4%), and the int8 model quantization with both int8 and int16 activations had the lowest post-conversion accuracies of 98.3% and 98.3%, respectively.

When we tested the model compression methods on our deployed iOS application, the fastest inference speed of 2.75 ms was achieved using float32 model compression running on the Core ML delegate (Table 7). For mobile CPU performance, int8 quantization with int16 model activations achieved the fastest inference time for both single and multi-threaded processing, with average speeds up to 9.20 ms and 5.56 ms, respectively (Tables 8 and 9). For mobile GPU performance, float32 model compression had the fastest inference time of 3.58 ms on the Metal delegate (Table 10). For performance on the Core ML delegate, float32 model compression had the fastest inference time on all mobile devices (2.75 ms).

**Table 7.** Inference times (ms) from the model conversion optimization methods that we studied and deployed on iPhone 11 and iPhone 13 running the Core ML delegate with parallel processing splits between CPU, GPU, and NPU computing. The fastest prediction times for each device are bolded.

| Device | Conversion 1 | Conversion 2 | Conversion 3 | Conversion 4 | Conversion 5 |
| --- | --- | --- | --- | --- | --- |
| iPhone 11 | **2.75ms** | 2.77ms | 2.82ms | 8.00ms | 10.47ms |
| iPhone 13 | **3.34ms** | 3.35ms | 3.51ms | 9.86ms | 11.21ms |

**Table 8.** Inference times (ms) from the model conversion optimization methods that we studied and deployed on iPhone 8+, iPhone X, iPhone 11, and iPhone 13 with single-threaded CPU computation. The fastest prediction times for each device are bolded.

| Device | Conversion 1 | Conversion 2 | Conversion 3 | Conversion 4 | Conversion 5 |
| --- | --- | --- | --- | --- | --- |
| iPhone 8+ | 23.01ms | 22.50ms | 22.87ms | **19.54ms** | 22.18ms |
| iPhone X | 21.14ms | 20.92ms | 20.92ms | **18.61ms** | 20.62ms |
| iPhone 11 | 17.04ms | 16.06ms | 16.38ms | **9.20ms** | 10.42ms |
| iPhone 13 | 11.26ms | 11.12ms | 11.20ms | **9.92ms** | 11.11ms |

**Table 9.** Inference times (ms) from the model conversion optimization methods that we studied and deployed on iPhone 8+, iPhone X, iPhone 11, and iPhone 13 with multi-threaded CPU computation. The fastest prediction times for each device are bolded.

| Device | Conversion 1 | Conversion 2 | Conversion 3 | Conversion 4 | Conversion 5 |
| --- | --- | --- | --- | --- | --- |
| iPhone 8+ | 14.81ms | 14.34ms | 14.83ms | **13.96ms** | 16.44ms |
| iPhone X | 13.96ms | 12.68ms | 14.51ms | **11.65ms** | 12.73ms |
| iPhone 11 | 10.30ms | 9.69ms | 9.45ms | **5.56ms** | 7.90ms |
| iPhone 13 | 7.05ms | 6.89ms | 6.87ms | **6.41ms** | 7.65ms |

**Table 10.** Inference times (ms) from the model conversion optimization methods that we studied and deployed on iPhone 8+, iPhone X, iPhone 11, and iPhone 13 with GPU computation using the Metal delegate. The fastest prediction times for each device are bolded.

| Device | Conversion 1 | Conversion 2 | Conversion 3 | Conversion 4 | Conversion 5 |
| --- | --- | --- | --- | --- | --- |
| iPhone 8+ | **24.03ms** | 24.54ms | 25.14ms | 25.31ms | 24.20ms |
| iPhone X | 20.90ms | 20.76ms | **20.61ms** | 22.53ms | 20.79ms |
| iPhone 11 | **10.28ms** | 10.35ms | 10.38ms | 11.53ms | 10.78ms |
| iPhone 13 | **3.58ms** | 3.60ms | 3.61ms | 4.13ms | 3.61ms |

The Core ML and Metal delegates (i.e., parallel CPU, GPU, and NPU processing and direct GPU processing) had the best performances on newer mobile devices like iPhone 11 and iPhone 13 with inference times of 2.75 ms and 3.58 ms, respectively. In contrast, CPU processing had slower inference times of 9.20 ms and 5.56 ms on single and multi-threaded CPUs. For older devices like the iPhone 8+ and iPhone X, multi-threaded CPU processing achieved faster inference times compared to single-threaded CPU and GPU processing.

## IV. Discussion

This study presents the development and mobile deployment of *StairNet*, a vision-based automated stair recognition system powered by deep learning. Building on *ExoNet* [20] – the largest and most diverse open-source dataset of wearable camera images of walking environments - we designed a new dataset specifically for stair recognition. We then developed a light- weight and efficient deep learning model, which accurately predicted the stair environments with 98.4% accuracy. We also studied the effects of different mobile devices (i.e., iPhone 8+, iPhone X, iPhone 11, and iPhone 13), model compression and optimization methods, and backend processing options, including single-threaded CPU, multi-threaded CPU, GPU, and a combination of CPU, GPU, and NPU in parallel. Of the systems that we tested, our highest performing model showed negligible reductions in prediction accuracy (<0.001%) due to the model conversion for mobile deployment and achieved an average inference time of 2.75 ms on an iPhone 11. In light of these results, the *StairNet* system has the ability to improve the autonomy and safety of human-robot locomotion via accurate and fast environment perception.

Compared to the original *ExoNet* studies [21]-[23], our system achieved significantly higher prediction accuracies (Table 11). This was accomplished by re-labelling the stair images in the *ExoNet* database using new class definitions to increase the precision of the cut-off points between different environmental states. Prior to our dataset revision, we developed a preliminary deep learning model, which gave low validation accuracies of 53.6% and 66.4% for the transition classes IS-LG and LG-IS, respectively, resembling the original *ExoNet* results [21]-[23]. We then retested the same model on our revised dataset, and the prediction accuracies increased to 84.4% and 87.5% for IS-LG and LG-IS, respectively, therein showing the benefit of our new class definitions. We used Google Cloud TPU accelerators to study the effects of different model and training parameters on performance, therein allowing for more extensive hyperparameter optimization and running our development pipeline in a fraction of the time compared to using NVIDIA graphics cards P100, T4, and V100.

**Table 11.** Review of vision-based stair recognition systems for wearable robotics. The dataset size (i.e., number of images) and test accuracy are only for the environment classes relating to level-ground walking and stair ascent. The systems are organized in terms of test accuracy (%).

| Reference | Camera | Position | Dataset Size | Classifier | Computing Device | Test Accuracy |
| --- | --- | --- | --- | --- | --- | --- |
| [7] | RGB | Waist | 7,284 | Convolutional neural network | NVIDIA Titan X | 99.6% |
| [8] | Depth | Chest | 170 | Heuristic thresholding and edge detector | Intel Core i5 | 98.8% |
| [10] | Depth | Leg | 8,455 | Support vector machine | Intel Core i7-2640M | 98.5% |
| **StairNet** | **RGB** | **Chest** | **515,452** | **Convolutional neural network** | **Google Cloud TPU** | **98.4%** |
| [13] | Depth | Leg | 3,000 | Convolutional neural network | NVIDIA Quadro P400 | 96.8% |
| [9] | Depth | Leg | 109,699 | Cubic kernel support vector machine | Intel Core i7-2640M | 95.6% |
| [6] | RGB | Chest | 34,254 | Convolutional neural network | NVIDIA TITAN Xp | 94.9% |
| [11] | RGB | Head | 123,979 | Bayesian deep neural network | NVIDIA Jetson TX2 | 93.2% |
| [12] | RGB | Leg | 123,954 | Bayesian deep neural network | NVIDIA Jetson TX2 | 92.4% |
| [21]–[23] | RGB | Chest | 542,868 | Convolutional neural network | Google Cloud TPU | 70.8% |

The *StairNet* system showed a number of benefits over other stair recognition systems for wearable robots (Table 11). Most systems use statistical pattern recognition and machine learning algorithms that require time-consuming and suboptimal hand engineering. In contrast, our deep learning model replaces these hand-designed features with multilayer networks that can automatically and efficiently learn the optimal image features from training data. However, this approach is susceptible to outliers and/or misclassified images in the training set, which were addressed through our dataset revision and quality control. We also developed *StairNet* using significantly more data than other CNN-based stair recognition systems. For example, Laschowski et al. [6] developed one of the first stair recognition systems using convolutional neural networks. However, their dataset included only ∼34,000 labelled images whereas *StairNet* has over 515,000 labelled images of indoor and outdoor walking environments throughout the summer, fall, and winter. These differences can have important practical implications since deep learning often requires significant and diverse training data to promote generalization [14].

There are also no explicit requirements for the pose of our wearable camera. This is one of the benefits of our system compared to other designs like [8], which relied on heuristics and meticulous rule-based thresholds for the dimensions of the user and environment. Our deep learning model appeared highly robust to changes in yaw, pitch, and roll of the camera during natural walking in unconstructed environments. There are also no explicit assumptions on the gait pattern. However, we did not conduct formal experiments to study the robustness of our perception system to persons of different walking abilities.

Another unique contribution of this study was deployment and inference on mobile devices. When deployed, *StairNet* achieved an inference time up to 2.75 ms. Other environment recognition systems for robotic prosthetic legs and exoskeletons have shown prediction speeds ranging from 0.9–50 ms [11], [12], [29]-[33], though many were tested on desktop computers. Of those deployed on mobile devices [11], [12], our system was ∼4.5x faster. To achieve fast inference speeds, we used an efficient and lightweight convolutional neural network designed for mobile and embedded vision applications. We then studied different model compression and optimization methods to reduce the model size and increase compatibility with the mobile hardware accelerators. We also studied our compressed deep learning models in a deployed environment using different inference delegates powered by CPU, GPU, and/or NPU backend computing to further accelerate the inference predictions, which is another novel contribution of our research.

Despite these advances, our study has several limitations. We estimated the classification accuracy during inference using the *StairNet* test set. While test sets are common in deep learning [14], our accuracy performance was not evaluated in a deployed environment where additional factors, such as camera obstructions, could impact accuracy. Also, the feasibility of our environment recognition system to support robot control was only considered from a theoretical perspective such that the speed and accuracy of the environment state predictions and their impact on user safety was not studied. Moreover, the *StairNet* system was mainly designed to support research and development. To improve useability and human-computer interaction, we plan to transition to a smart glasses design. Also, limited comparisons could be made between the performance and efficiency of our system and others since we used different image datasets for training and testing. We hope to address this limitation by making the *StairNet* dataset publicly available to the research community.

It is important to emphasize that our perception systems are meant to supplement, not replace, the existing intent recognition systems that use mechanical, inertial, and/or surface electromyographic data to estimate the current state of the humanrobot-environment system [1]. We view computer vision as a means to improve the speed and accuracy of locomotion mode (intent) recognition by minimizing the search space of potential solutions based on the perceived walking environment. Accordingly, sensor fusion methods will need to be explored in the future to integrate computer vision with the existing intent recognition systems used for high-level control of robotic prosthetic legs and exoskeletons.

## V. Conclusion

The high accuracy (98.4%) and low latency (2.75ms) of our new AI-powered stair recognition system deployed on mobile devices shows the feasibility to develop real-world computer vision systems for autonomous control and planning of robotic prosthetic legs, exoskeletons, and other assistive technologies for human locomotion. Moving forward, we plan to transition to a smart glasses design in order to improve human-computer interaction and integrate our perception system into the robot control system for environment-adaptive locomotion mode recognition.

## Acknowledgment

We thank the University of Toronto MADLab for their help with the mobile software deployment and Shehryar Saharan for Fig. 1. This research is dedicated to the people of Ukraine in response to the 2022 Russian invasion.

